# Weberviruses are gut-associated phages that infect *Klebsiella* spp

**DOI:** 10.1101/2024.11.04.620279

**Authors:** Samuel J. T. Dawson, Preetha Shibu, Sara Garnett, Fiona Newberry, Thomas C. Brook, Tobi Tijani, Magdalena Kujawska, Lindsay J. Hall, Anne L. McCartney, David Negus, Lesley Hoyles

## Abstract

Weberviruses are bacteriophages (phages) that can infect and lyse clinically relevant, multidrug-resistant (MDR) strains of *Klebsiella*. They are an attractive therapeutic option to tackle *Klebsiella* infections due to their high burst sizes, long shelf life and associated depolymerases. In this study we isolated and characterized seven new lytic phages and compared their genomes with those of their closest relatives. Gene-sharing network, ViPTree proteome and *terL* gene-sequence-based analyses incorporating all publicly available webervirus genomes [*n*=258 from isolates, *n*=65 from metagenome-assembled genome (MAG) datasets] confirmed the seven phages as members of the genus *Webervirus* and identified a novel genus (*Defiantjazzvirus*) within the family *Drexlerviridae*. Using our curated database of 265 isolated phage genomes and 65 MAGs (*n*=330 total), we found that weberviruses are distributed globally and primarily associated with samples originating from the gut: sewage (154/330, 47 %), wastewater (83/330, 25 %) and human faeces (66/330, 20 %). We identified three distinct clusters of potential depolymerases encoded within the 330 genomes. Due to their global distribution, frequency of isolation and lytic activity against the MDR clinical *Klebsiella* strains used in this study, we conclude that weberviruses and their depolymerases show promise for development as therapeutic agents against *Klebsiella* spp.

## INTRODUCTION

Members of the *Klebsiella pneumoniae* species complex are opportunistic pathogens that can cause serious hospital-acquired infections, and are major contributors to global deaths associated with antimicrobial resistance (Antimicrobial Resistance Collaborators, 2022). Carbapenem-resistant isolates of *K. pneumoniae* are resistant to a range of frontline β-lactam antibiotics. The difficulty of treating infections caused by such isolates with conventional antibiotics has resulted in the investigation of new therapeutic modalities, including bacteriophages (phages) and their gene products (Herridge et al., 2020). To realize the potential of phage therapy, it is important to comprehensively characterize phages with clinical use. Previously we isolated *Webervirus KLPN1* from the caecum of a healthy female, along with its host *K. pneumoniae* subsp. *pneumoniae* L4-FAA5 (Hoyles et al., 2015). In the current study, we successfully identified seven new representatives of the genus *Webervirus* using L4-FAA5 and multidrug-resistant (MDR) clinical isolates of *Klebsiella* spp. as isolation hosts. These hosts included *K. pneumoniae* PS_misc6, which encodes the carbapenem-degrading metallo-β-lactamase NDM, and *Klebsiella variicola* PS_misc5, a carbapenem-resistant clinical isolate that encodes the class D β-lactamase OXA-48 (Shibu, 2019).

As of 19 January 2025, the genus *Webervirus* encompassed 100 different phage species [<u>International Committee on Taxonomy of Viruses</u> (ICTV)]. With the exception of *Webervirus BUCT705* (isolated on *Stenotrophomonas maltophila*), all weberviruses described to date have been isolated on *Klebsiella* hosts, and have proven easy to recover from sewage, wastewater and, occasionally, intestinal contents (Herridge et al., 2020). Although *Webervirus F20* was originally described as being isolated on *Enterobacter aerogenes* (Mishra et al., 2012), this bacterium has subsequently been reclassified as *Klebsiella aerogenes*. Their high burst sizes (approximately 80 pfu/cell with a reported range between 27 and 142 pfu/cell) (Fang et al., 2022; Gilcrease et al., 2023; P. Li et al., 2024; Senhaji-Kacha et al., 2024; Ziller et al., 2024; Zurabov and Zhilenkov, 2021) and long shelf life make weberviruses ideal phages to work with for biotechnological and clinical applications (Fang et al., 2022; Herridge et al., 2020).

Currently, more than 130 different capsule types (K types) have been identified for *K. pneumoniae* by genetic analysis (Follador et al., 2016). Specific *K. pneumoniae* capsule types are strongly associated with virulence. For example, hypervirulent *K. pneumoniae* isolates are typically associated with capsule types K1, K2, K16, K28, K57 and K63 (Kabha et al., 1995; Lee et al., 2016; Marr and Russo, 2019; Mizuta et al., 1983; Yu et al., 2008). Additionally, capsule production by *Klebsiella* spp. has been implicated in protection from complement-mediated lysis and is recognized to play an important role in biofilm formation (Alvarez et al., 2000; Jensen et al., 2020). Capsule type has also been shown to be a major determinant of host tropism in *Klebsiella* phages (Beamud et al., 2023). Bacterial capsules are known to prevent phage attachment by masking cell-surface-associated receptor proteins (Dunstan et al., 2021; Scholl et al., 2005). To overcome this physical barrier, phages encode enzymes – frequently referred to as depolymerases – that selectively degrade polysaccharides or polypeptides that comprise the bacterial capsule (Cai et al., 2023; Dunstan et al., 2021; Hoyles et al., 2015; Majkowska-Skrobek et al., 2016; Pertics et al., 2021). Weberviruses tend to have narrow host ranges (Hoyles et al., 2015; Pertics et al., 2021). However, our previous (and ongoing) work has suggested that their depolymerases can degrade the capsules of non-host *Klebsiella* spp. (Hoyles et al., 2015). Depolymerase activity is common to weberviruses, and is being actively investigated as a tool to hydrolyse capsules of *Klebsiella* spp. that often hinder or make treatment with antimicrobials difficult (Cai et al., 2019; Majkowska-Skrobek et al., 2016; Pertics et al., 2021). For example, the webervirus depolymerase Depo32 has been shown to protect mice from otherwise lethal *K. pneumoniae* infections in a mouse model of disease (Cai et al., 2023). In addition, a webervirus (P39) has recently been used in combination with another lytic phage (P24, *Przondovirus*) to decolonize mice of carbapenem-resistant *K. pneumoniae* (Fang et al., 2022).

Here we describe our new webervirus phages and their lytic and depolymerase activities against clinical MDR *Klebsiella* spp., and compare their genomes with those of their closest relatives. The increased ease with which metagenome-associated viruses can be interrogated via PhageClouds (Rangel-Pineros et al., 2021) and NCBI also led us to determine whether weberviruses are readily detectable within recent metagenome-derived phage datasets.

## METHODS

### Bacterial strains

Details of all *Klebsiella* strains included in this study are given in **Table 1**. The antimicrobial resistance profiles of the isolates, determined according to EUCAST guidelines as described previously (Shibu et al., 2021), can be found in **Supplementary Table 1**.

**Table 1.**
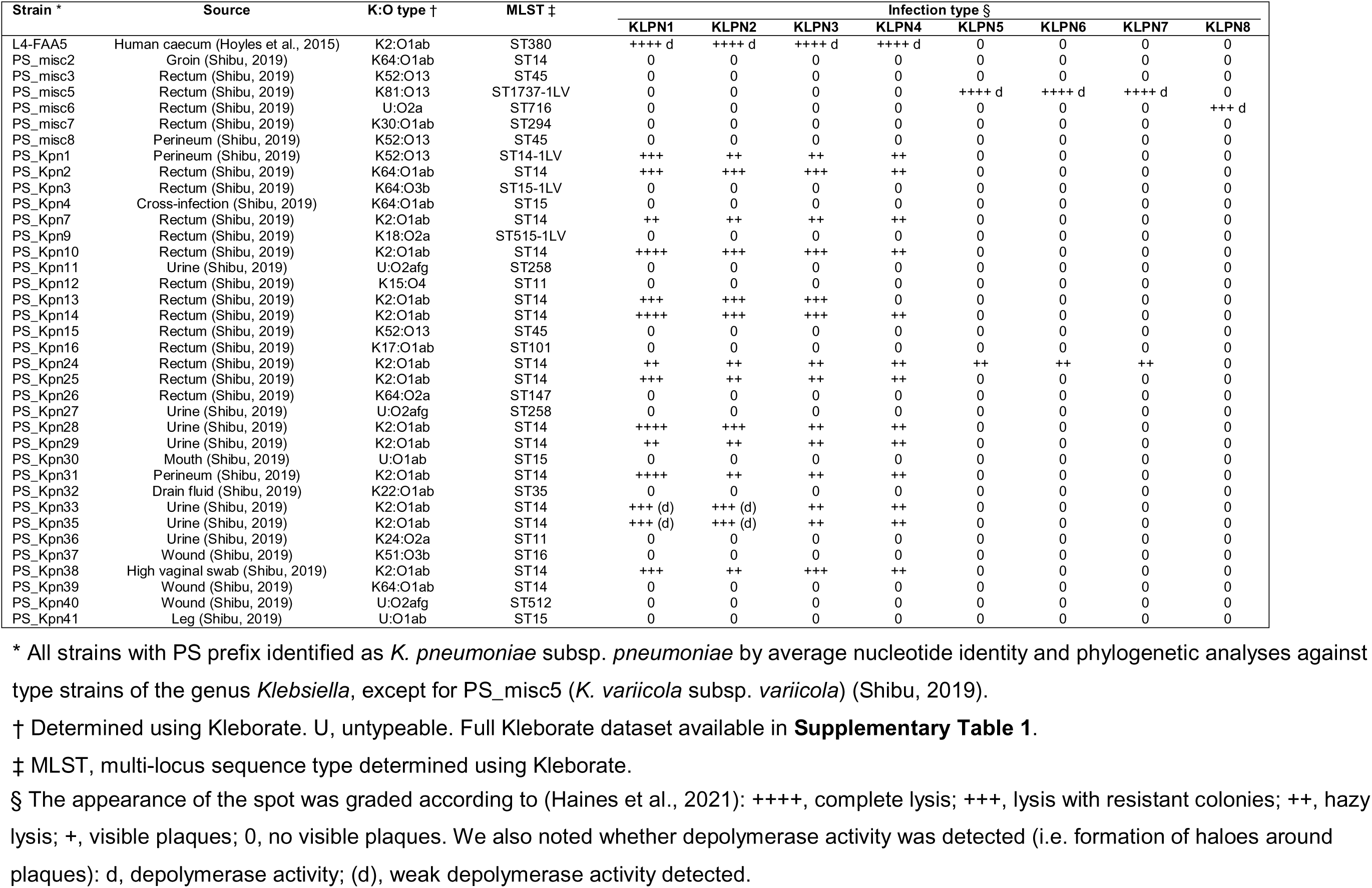
Strains of *Klebsiella* included in this study and their phage infection profiles.

### Generation of sequence data for bacterial isolates

Genomes for clinical strains included in this study were generated as described previously (Shibu et al., 2021). Illumina and Oxford Nanopore Technologies sequence data for *K. pneumoniae* L4-FAA5 were generated by microbesNG (Birmingham, UK) as described previously (Newberry et al., 2023). CheckM2 v0.1.3 (Chklovski et al., 2022) was used to confirm the quality (in terms of completeness and contamination) of all assembled genomes. Kleborate v3.1.2 (Lam et al., 2021; Wyres et al., 2016) was used to assign sequence types (STs), and capsule (K) and lipopolysaccharide (O) types to genomes.

### Isolation and propagation of phages

Filter-sterilized sewage samples were screened for phages as described previously (Smith-Zaitlik et al., 2022) using *Klebsiella* strain L4-FAA5, PS_misc5 or PS_misc6 as inoculum (**Table 1**). Pure phage stocks were prepared from phage-positive samples as described previously (Hoyles et al., 2015).

### Isolation of phage DNA

Phages vB_KpnS-KLPN2, vB_KpnS-KLPN3 and vB_KpnS-KLPN4 were precipitated from 100 ml of each lysate as described previously (Hoyles et al., 2015).Phages vB_KvaS-KLPN5, vB_KvaS-KLPN6, vB_KvaS-KLPN7 and vB_KpnS-KLPN8 were concentrated and DNA extracted as described previously (Smith-Zaitlik et al., 2022).

### Transmission electron microscopy

Transmission electron micrographs (TEMs) of phages vB_KpnS-KLPN2, vB_KpnS-KLPN3 and vB_KpnS-KLPN4 were generated as described previously (Hoyles et al., 2015). TEMs for phages vB_KvaS-KLPN5, vB_KvaS-KLPN6, vB_KvaS-KLPN7 and vB_KpnS-KLPN8 were generated and analysed as described previously (Smith-Zaitlik et al., 2022).

### Phage genome sequencing, assembly and annotation

Assembled genomes (from Illumina short-read sequences) for phages vB_KpnS-KLPN2, vB_KpnS-KLPN3 and vB_KpnS-KLPN4 were generated by microbesNG (Shibu et al., 2021). For phages vB_KvaS-KLPN5, vB_KvaS-KLPN6, vB_KvaS-KLPN7 and vB_KpnS-KLPN8, sequence data were generated on an Illumina MiSeq at Nottingham Trent University (Smith-Zaitlik et al., 2022). Quality of raw sequence data was assessed using FastQC v0.11.9. Reads had a mean phred score above 30 and no adapter contamination, so data were not trimmed.

All genomes were assembled using SPAdes v3.13.0 (default settings) (Bankevich et al., 2012), and visualized to confirm circularization of genomes using Bandage v0.8.1 (Wick et al., 2015). CheckV v1.0.1 (checkv-db-v1.5; (Nayfach et al., 2021a)) was used to determine contamination and completeness of the genomes. Genes in all phage genomes included in this study (**Supplementary Table 2**) were predicted and annotated using Pharokka v1.6.1 (v1.4.0 databases) (Bouras et al., 2023).

### Comparison of webervirus genomes

ViPTree v4.0 (Nishimura et al., 2017) was used to determine whether the seven phage genomes were closely related to previously described double-stranded DNA viruses. Based on our initial findings (not shown) we curated a list of all known webervirus sequences available from NCBI GenBank on 19 January 2025. We also identified unclassified weberviruses and closely related phage in NCBI using the INPHARED database (1Jan2025 dataset; (Cook et al., 2021)) and vConTACT v2.0 (**Supplementary Table 2**).

### Identification of weberviruses in metagenomic datasets

We used PhageClouds (Rangel-Pineros et al., 2021) to identify relatives of weberviruses in metagenome-assembled genome (MAG) datasets. PhageClouds is an online resource that allows researchers to search a reference dataset of ∼640,000 phage genomes for phages with genomes related to query sequences. The genome of *Webervirus KLPN1* was searched against the PhageClouds database with a threshold of 0.15, as we had previously looked for relatives of this phage in metagenomic datasets and are interested in gut-associated phage communities (Hoyles et al., 2015). The nucleotide sequences of the relevant phage MAGs (from (Camarillo-Guerrero et al., 2021; Gregory et al., 2020; Tisza and Buck, 2021)) were recovered from the relevant datasets.

Additional webervirus MAGs were identified using a search of the NCBI nucleotide database for Bacteriophage sp. [search term: (txid38018) AND MAG]; the sequences (*n*=8138) were filtered for genomes of between 30 Kbp and 60 Kbp in length (*n*=2540): genes were predicted using Prodigal v.2.6.3 (Hyatt et al., 2010) and the proteomes added to the INPHARED database and analysed using vCONTact2. CheckV was used (as described above) to determine contamination and completeness of the MAG dataset (**Supplementary Table 3**).

The MAG sequences were analysed using ViPTree v4.0 to confirm their affiliation with the genus *Webervirus*. They were also annotated with Pharokka and included in a vConTACT2 analysis with our curated set of webervirus genomes. The genomes of all weberviruses were compared with one another using taxmyPHAGE v0.3.3, which uses a Python implementation of the VIRIDIC algorithm to calculate intergenomic genomic similarities (Millard et al., 2024). The matrix created from the similarity values was visualized using tidyheatmaps v0.2.1 (Mangiola and Papenfuss, 2020).

### Phylogenetic relationships among weberviruses

Nucleotide sequences of the large-subunit terminase (*terL*) genes, predicted by Pharokka, were used to create a multiple-sequence alignment (Clustal Omega 1.2.2 implemented in Geneious Prime v2024.0.5; options – group sequences by similarity, 5 representative iterations). This alignment was used to create a bootstrapped (100 replicates) maximum-likelihood tree (PhyML v3.3.20180621, JC69 algorithm).

### Distribution of weberviruses

The distribution of weberviruses was determined by identifying the source and geographical location information for the GenBank genomes (including our seven new genomes; **Supplementary Table 2**) and the MAGs (**Supplementary Table 3**). Data were aggregated based on isolation source or geographical location, with these latter data visualized using the R package rworldmap v1.3.8 (South, 2011).

### Host prediction for MAGs

The CRISPR Spacer Database and Exploration Tool (Dion et al., 2021) and HostPhinder 1.1 (Villarroel et al., 2016) were used to predict hosts for the weberviruses recovered from metagenomic datasets. MAGs were also subject to a BLASTN search against the Unified Human Gastrointestinal Genome (UHGG) CRISPR spacer database according to (Nayfach et al., 2021b). For this, a BLASTN database was created from 1,846,441 spacers from 145,053 CRISPR arrays from 79,735 UHGG genomes (Nayfach et al., 2021b). Spacers were searched against viral genomes using BLASTN from the blast+ package v.2.12.0 (options:-dust=no; -word-size=18); a maximum of one mismatch or gap was allowed over ≥95 % of the spacer length. iPHoP v1.3.3 (Roux et al., 2023), an automated command-line pipeline for predicting host genus of novel bacteriophages and archaeoviruses based on their genome sequences, was also used to analyse the MAGs.

### Identification of potential depolymerases among weberviruses

A BLASTP database was created using amino acid sequences from experimentally validated webervirus depolymerases and a BLASTP search was ran versus all webervirus genomes. Sequences used to build the BLAST database are available from figshare (doi:10.6084/m9.figshare.28603070). Clustal Omega v1.2.2 alignments were created in Geneious Prime (default settings; 2024.0.5). RAxML v 8.2.11 (-m PROTGAMMABLOSUM62 -f a -x 1 -N 100 -p 1) was used to generate a bootstrapped (100 replicates) maximum-likelihood tree from the multiple-sequence alignment.

## RESULTS

### Seven new weberviruses lyse a range of clinically relevant *Klebsiella* spp

Seven phages were isolated on two different strains of *K. pneumoniae* subsp. *pneumoniae* (L4-FAA5 – vB_KpnS-KLPN2, vB_KpnS-KLPN3, vB_KpnS-KLPN4; PS_misc6 – vB_KpnS-KLPN8) and one strain of *K. variicola* subsp. *variicola* (PS_misc5 – vB_KvaS-KLPN5, vB_KvaS-KLPN6, vB_KvaS-KLPN7). All our sewage samples yielded *Klebsiella*-infecting phages. Strain L4-FAA5 (K2:O1ab, ST380) was originally isolated from human caecal effluent along with *Webervirus KLPN1* (Hoyles et al., 2015), while strains PS_misc5 (K81:O13, ST1737-1LV) and PS_misc6 (untypeable:O2a, ST716) were part of a collection (*n*=36) of clinical MDR and/or carbapenem-resistant *Klebsiella* isolates currently being used in our laboratory in phage-related and other studies (Shibu, 2019) (**Supplementary Table 1**).

TEM showed the seven phages had a mean capsid diameter of 57.5 nm and a mean tail length of 157.5 nm (**Supplementary Figure A**). Host-range analysis showed the seven phages had different infection profiles (**Table 1**). KLPN1, our original webervirus isolated on *K. pneumoniae* L4-FAA5 (Hoyles et al., 2015), was included in analyses for comparative purposes. KLPN1, vB_KpnS-KLPN2, vB_KpnS-KLPN3 and vB_KpnS-KLPN4 completely lysed some, but not all, clinical isolates of *K. pneumoniae* with capsule:O antigen types K52:O13 and K64:O1ab. K2:O1ab isolates alone were infected by KLPN1 and vB_KpnS-KLPN2 to vB_KpnS-KLPN4, though vB_KpnS-KLPN4 was unable to infect one of the K2:O1ab strains (PS_Kpn13). Only on strain L4-FAA5 (K2:O1ab), isolated from human caecal effluent, was strong depolymerase activity observed with phages KLPN1 and vB_KpnS-KLPN2. Hazy lysis of strain PS_Kpn24 (K2:O1ab) was observed with phages vB_KvaS-KLPN5, vB_KvaS-KLPN6, vB_KvaS-KLPN7. Phages vB_KvaS-KLPN5 to vB_KvaS-KLPN7 showed strong lytic and depolymerase activity on *K. variicola* PS_misc5 (K81:O13) alone, while vB_KpnS-KLPN8 lysed *K. pneumoniae* PS_misc6 (untypeable capsule:O2a) with depolymerase activity on this host.

### Genome-based analyses of publicly available sequence data triples the number of authenticated webervirus genomes

Bandage (data not shown) and CheckV (**Supplementary Table 2**) analyses confirmed the genomes of vB_KpnS-KLPN2, vB_KpnS-KLPN3, vB_KpnS-KLPN4, vB_KvaS-KLPN5, vB_KvaS-KLPN6, vB_KvaS-KLPN7 and vB_KpnS-KLPN8 were circular and complete. None of the genomes was contaminated. An initial online ViPTree analysis showed vB_KpnS-KLPN2, vB_KpnS-KLPN3, vB_KpnS-KLPN4, vB_KvaS-KLPN5, vB_KvaS-KLPN6, vB_KvaS-KLPN7 and vB_KpnS-KLPN8 belonged to the genus *Webervirus* (data not shown). All publicly available webervirus genomes (available as of 19 January 2025) were downloaded from GenBank to allow comparison with our newly sequenced phages, and for inclusion in the INPHARED vCONTact2 database if not already included in the 1Jan2025 release. Among the other 264 genomes from phage isolates included in this study, 226 were of high quality, 43 were complete and two were of medium quality; none of these genomes was contaminated.

In addition, we used PhageClouds to identify potential webervirus MAGs. Fifty-four of the PhageClouds hits represented MAGs derived from the Gut Phage Database (GPD) (Camarillo-Guerrero et al., 2021), six were from the Cenote Human Virome Database (CHVD) (Tisza and Buck, 2021) and two were from the Gut Virome Database (GVD) (Gregory et al., 2020). MAG Ma_2019_SRR413710_NODE_378_length_50715_cov_48.086538 from the GVD was identical to uvig_330395 from the GPD (Camarillo-Guerrero et al., 2021) so was removed from further analyses (PhageClouds scores identical, 100 % pairwise identity as assessed using VIRIDIC; an unsurprising finding as both MAGs are derived from the same dataset (Ma et al., 2018)). Similarly, two MAGs from the GPD were also found to be identical: uvig_314355 and uvig_315584 were high-quality genomes both derived from the same four samples (SRR1952259, SRR1162648, SRR1162662, SRR1162654 (Tisza and Buck, 2021)); only uvig_314355 was retained for further analyses. Our inclusion of NCBI genomes listed as Bacteriophage sp. in a vCONTact2 analysis with the INPHARED database identified a further five potential webervirus MAGs recovered from faecal samples in Japan (Nishijima et al., 2022). In total, our dataset included 65 MAGs. The MAGs ranged from 10,230 to 55,276 nt (mean 42,392 nt) in length (**Supplementary Table 3**). Forty-seven of the 65 MAGs were determined to be complete or of high-quality (CheckV). Eight were of medium-quality and 10 were low quality, representing genome fragments (**Supplementary Table 3**). None of the MAGs was contaminated.

In addition to the 100 recognized weberviruses included in the ICTV and our seven new weberviruses, we identified 158 more webervirus genomes in NCBI and 65 webervirus MAGs. The 265 weberviruses isolated on bacteria mostly infected *K. pneumoniae* (**Supplementary Figure B**). A ViPTree analysis confirmed the affiliation of our 330 genomes with the genus *Webervirus* (**Figure 1**). The webervirus genomes often clustered based on geographical origin, irrespective of whether they came from phage isolates or MAGs (**Figure 1**).

**Figure 1.**
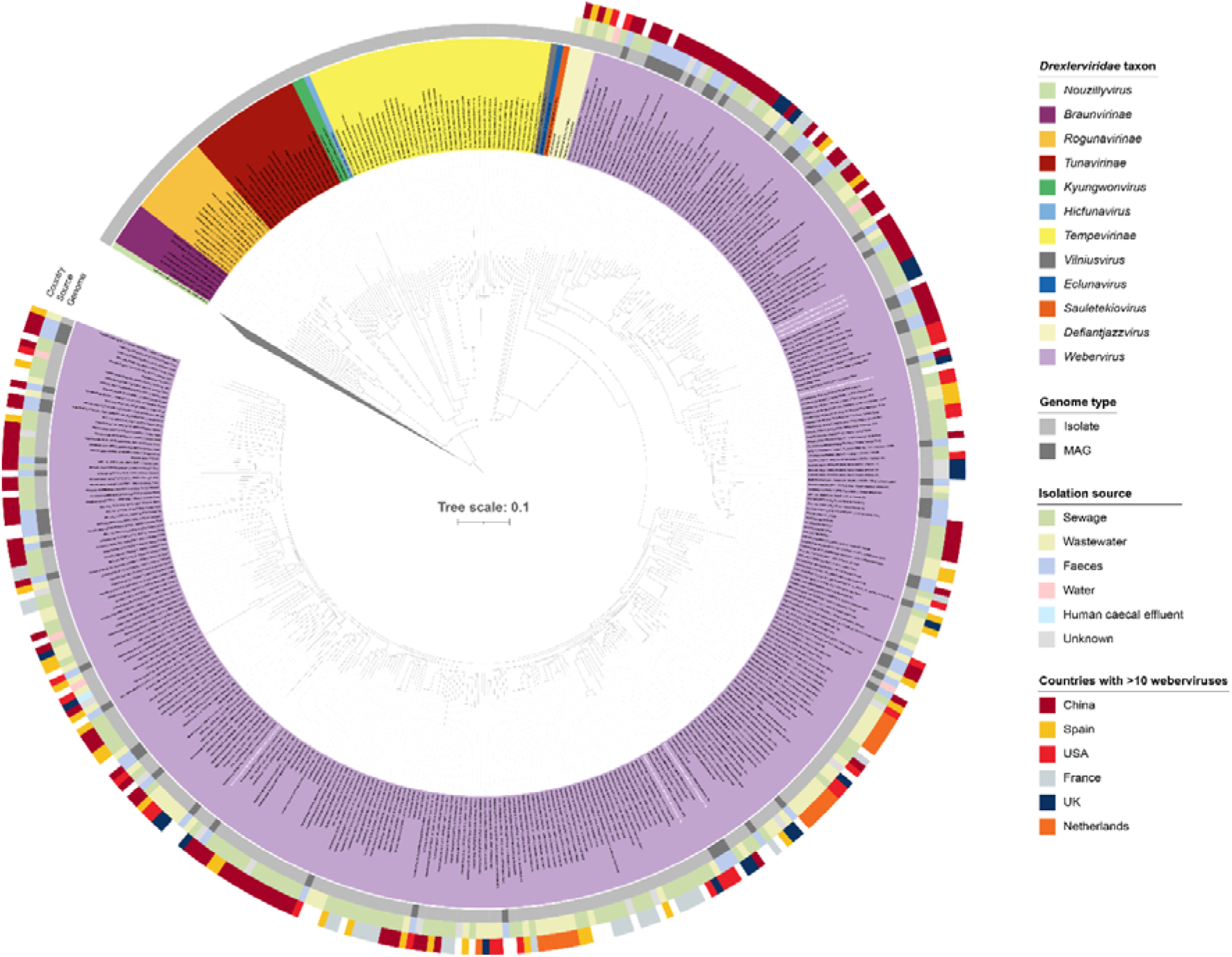
ViPTree-generated phylogenetic analysis of the family *Drexlerviridae*. The genus *Webervirus* is represented by 330 genomes. The names of our seven newly identified weberviruses are shown in white bold text. A potentially novel genus (*Defiantjazzvirus*) was identified during the curation of our dataset. The colours covering the virus names represent taxa within the family *Drexlerviridae*; the outgroup has been collapsed to aid visualization. The tree (ViPTree bionj) was rooted at the midpoint.

The monophyletic nature of the genus *Webervirus* was confirmed by phylogenetic analysis of *terL* gene sequences (99 % bootstrap support; **Figure 2a**). A gene-sharing network was created with all webervirus genomes included in this study (**Table 2**, **Table 3**) using vConTACT v2.0 (Bin Jang et al., 2019; Bolduc et al., 2017) and the INPHARED database (**Supplementary Figure C**). The network was filtered based on first and second neighbours of webervirus proteomes (**Supplementary Figure C**, **Figure 2b**). The vConTACT-based analysis confirmed findings from the ViPTree- and *terL*-based analyses with respect to affiliation of weberviruses included in this study.

**Figure 2.**
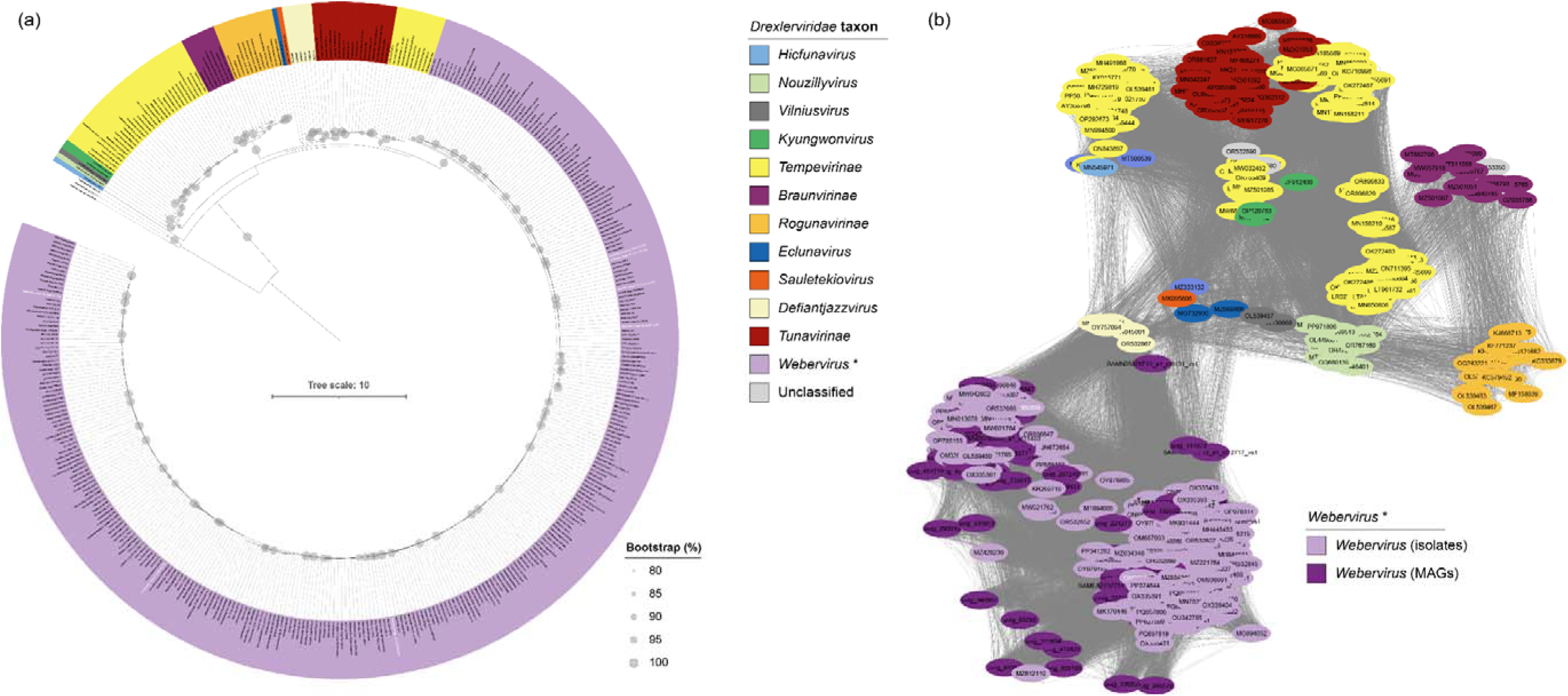
Further analyses of *Drexlerviridae* sequence data. (a) Phylogenetic relationships (maximum-likelihood tree) of members of the family *Drexlerviridae* based on analysis of large-subunit terminase (*terL*) nucleotide sequences encoded in phage genomes. Bootstrap values are expressed as a percentage of 100 replications; scale bar, mean number of nucleotide substitutions per position; the tree is rooted at the midpoint. (b) Gene-network-based analysis of proteomic data for members of the genus *Webervirus* and their nearest relatives. Full network shown in **Supplementary Figure C**. (a, b) The legend shown applies to both figures, with isolate and MAG proteomes differentiated in (b). Names of our seven newly identified weberviruses are shown in bold white text.

VIRIDIC analysis split the weberviruses into eight different clusters at the genus level, with most weberviruses affiliated with Cluster 1 (**Supplementary Figure D**, **Supplementary Table 4**). Clusters 3 (uvig_338855, uvig_63295), 4 (uvig_346479, uvig_474523), 5 (SAMN05826713_a1_ct6131_vs1), 6 (uvig_63387), 7 (uvig_340901), 8 (uvig_334913) and 9 (SAMN05826713_a1_ct12717_vs1) were all associated with low-quality MAGs (**Supplementary Table 3**). MAGs in these clusters shared <70 % identity with Cluster 1 phages (isolate and MAG genomes). The only other low-quality MAG included in the analysis (uvig_311634) was affiliated with Cluster 1 phages, sharing 33–72 % identity with them and highest similarity with a MAG (uvig_141073) in this cluster (**Supplementary Table 4**).

### Identification of a novel genus within the family *Drexlerviridae*

Our ViPTree analysis also identified a potential novel genus (referred to as *Defiantjazzvirus*) comprising six representatives within the family *Drexleviridae* and closely related to the genus *Webervirus* (**Figure 1**). Analysis of *terL* gene sequences showed this genus to be monophyletic (97 % bootstrap support; **Figure 2a**). vConTACT-based analysis demonstrated that the six genomes associated with *Defiantjazzvirus* clustered together but separately from all other phage groups included in the analysis (**Figure 2b**). VIRIDIC analysis showed defiantjazzvirus genomes to share 81–97 % genome identity with one another, and 27–42 % identity with members of the genus *Webervirus* (**Supplementary Figure D**, **Supplementary Table 4**). Based on current recommendations, the six genomes (sharing >70 % nucleotide identity across their full-length genomes) represent a novel genus comprising five species (**Supplementary Table 4**) (Turner et al., 2021). Comparison of the defiantjazzvirus genomes with non-webervirus *Drexleviridae* genomes confirmed the genus *Defiantjazzvirus* represents a novel genus within the family *Drexleviridae*, with the six defiantjazzvirus genomes sharing between 0.2 and 35 % genome identity with their closest non-webervirus relatives (**Supplementary Table 5**). Representatives of the genus *Defiantjazzvirus* infect *K. pneumoniae*, *K. michiganensis* and *K. oxytoca* (**Supplementary Table 2**).

### Webervirus MAGs are predicted to infect *Klebsiella*

To confirm the MAGs were weberviruses which infected *Klebsiella* spp. we attempted to predict their bacterial hosts. CRISPR spacers can be used to predict hosts of unknown phages, as spacers represent biological records of past phage–bacteria interactions. Each of the seven new phage genomes (nt sequences) we generated was uploaded to CRISPR Spacer Database and Exploration Tool (Dion et al., 2021). None of the phages could be assigned to known hosts using this tool. Using the BLASTN approach of (Nayfach et al., 2021b) with the MAG sequences, only SAMEA2737751_a1_ct5309 had sufficient coverage; this MAG had two hits to *Klebsiella* species (*K. pneumoniae* and *K. variicola*). iPHoP predicted hosts for 84/330 of the webervirus genomes included in this study; *Escherichia* was predicted to be the host for 21 of the MAGs and 59 of the isolates at the genome and genus levels (**Supplementary Table 6**). Only NC_049845.1, OR532813.1, OR532891, PQ337355 and PQ519586 – all representing isolated phages (**Supplementary Table 2**) – were predicted to have a *Klebsiella* host at the genus level. HostPhinder 1.1 (Villarroel et al., 2016) was able to predict hosts for our KLPN phages, with all assigned to *Klebsiella pneumoniae*. Consequently, this tool was used to predict hosts for the *Webervirus* MAGs (**Supplementary Table 7**). All were predicted to infect *Klebsiella*.

### Depolymerases are readily detected in webervirus genomes

As our newly isolated phages all displayed apparent depolymerase activity against one or more hosts, we aimed to identify potential depolymerases encoded within the genomes of weberviruses. Detection and characterization of these enzymes may identify standalone therapeutics or help inform on host tropism. Currently, four experimentally validated depolymerases from weberviruses have been reported in the literature: depoKP36 (Majkowska-Skrobek et al., 2016), Depo32 (Cai et al., 2023), DpK2 (Dunstan et al., 2021) and B1dep (Pertics et al., 2021). These four depolymerases were used to create a BLASTP database to interrogate the 330 webervirus genomes for similar amino acid sequences.

Using thresholds of >50 % coverage, >50 % identity and sequence length >800 aa, 33/330 webervirus proteomes returned hits against the validated depolymerases (**Figure 4**; **Supplementary Table 8**). Phylogenetic analysis and amino acid identity values revealed that the depolymerases clustered into three distinct groups, each with high bootstrap support (85–100 %; **Figure 4**). Group 1 comprised four sequences and did not contain an experimentally validated depolymerase sequence. Group 2 contained four sequences including the functionally characterized depolymerase depoKP36. Group 3 contained most the sequences (26/33 predicted depolymerases) and included the characterized depolymerases DpK2, Depo32 and B1dep, and depolymerases encoded by four MAGs. Sequences belonging to Group 3 had a high level of conservation as indicated by short branch lengths and sequence alignments (**Supplementary Figure E**). Amino acid alignment of all 33 predicted depolymerases also revealed a high level of N-terminal sequence conservation.

## DISCUSSION

### Isolation and characterization of seven new weberviruses

Studies from a diverse range of geographical locations have reported the isolation or detection of weberviruses from samples associated with the human gut (e.g. wastewater, sewage, faeces, caecal effluent) (Herridge et al., 2020). To date, the majority of weberviruses have been isolated using *K. pneumoniae* as a host (**Supplementary Figure B**). However, weberviruses have been reported to infect other *Klebsiella* spp. including *K. oxytoca* (Brown et al., 2017; Park et al., 2017) and *K. aerogenes* (Hudson et al., 2021). In the present study, we isolated seven new weberviruses from sewage samples, including three phages (vB_KvaS-KLPN5, vB_KvaS-KLPN6, vB_KvaS-KLPN7) that were isolated using a strain of *K. variicola* as the host (**Figure 1**, **Figure 2**, **Supplementary Figure A**). To our knowledge, this is the first report of weberviruses infecting *K. variicola*, a recognized emerging human pathogen (Rodríguez-Medina et al., 2019) increasingly associated with carbapenem and colistin resistance (Kim et al., 2023; L. Li et al., 2024).

As the majority of the *Klebisella* spp. sensitive to lysis by our webervirsues are MDR strains, the lytic phages isolated as part of this study represent attractive future therapeutics for the treatment of drug-resistant isolates belonging to the *K. pneumoniae* species complex.

In agreement with previous work (Hoyles et al., 2015; Pertics et al., 2021), the weberviruses described herein exhibited relatively narrow host ranges when screened against a panel of *Klebsiella* (including 36 clinical MDR) isolates representing a range of STs and capsule (K) types (**Table 1**). Phage host range is very much related to isolation host rather than phage phylogeny, with lysis appearing to be restricted based on K type. Phage-encoded depolymerases, therefore, contribute to host tropism and previous studies have identified that weberviruses encode functionally active depolymerases (Cai et al., 2023; Dunstan et al., 2021; Majkowska-Skrobek et al., 2016; Pertics et al., 2021). While performing our host-range analysis, we observed the presence of haloes indicative of depolymerase activity for a small number of phage–host combinations and we, therefore, undertook a bioinformatic analysis (**Figure 4**) to identify potential depolymerase enzymes encoded within webervirus genomes. Our BLASTP search identified 33 potential depolymerases which clustered into three distinct groups. The lack of an experimentally validated depolymerase sequence in Group 1 and the overall low amino acid identity shared with characterized webervirus depolymerases (< 21 %) makes it difficult to draw conclusions related to the biological activity of these four proteins. Sequences OP978314.1_CDS_0059 and OP978315.1_CDS_0001 belong to a phage, and its evolved variant, respectively, which were characterized as part of the same study in Australia (Ngiam et al., 2024). These phages were propagated on *K. pneumoniae* 52L145 (K2:O1). According to NCBI, the isolation host of phage OP413832.1, which encodes predicted depolymerase OP413832.1_CDS_0043, is *K. pneumoniae* BS317-1 (K57:O1) (assembly accession GCF_015290145.1). No information is available for the isolation host of the phage OR532859.1 which encoded the remaining predicted Group 1 depolymerase. These data suggest that, if active, Group 1 depolymerases may hydrolyse K2 and/or K57 capsules. However, experimental validation is required.

Group 2 depolymerases are likely to be hydrolyse the K63 capsule as these sequences clustered with the experimentally validated depolymerase depoKP36, previously shown to degrade the K63 capsule of *K. pneumoniae* (Majkowska-Skrobek et al., 2016). Group 3 contained the majority of the predicted depolymerases and all shared high sequence similarity with the webervirus depolymerases Depo32, DpK2 and B1dep (**Supplementary Table 8**). These enzymes have been shown to selectively degrade the *K. pneumoniae* K2 capsule (Cai et al., 2023; Dunstan et al., 2021; Pertics et al., 2021) and are highly likely to be specific for this capsule type. The high level of sequence identity observed at the N-terminal of all the identified depolymerases is likely due to this region being responsible for anchoring the baseplate of the phage virion, and as such it is often highly conserved (Knecht et al., 2019; Latka et al., 2019). Structural analysis of Depo32 from phage GH-K3 has revealed that, in addition to the N-terminal domain, Depo32 contains a short neck helix and connection domain (residues 186–271), a β-helix domain (residues 272–642), a connection helix domain (residues 643–666), a carbohydrate-binding module (residues 667–846), and a C-terminal domain (residues 847–907) (Cai et al., 2023). It is the β-helix domain that is responsible for hydrolysis of the polysaccharide capsule. Given the high level of amino acid identity between Depo32 and the amino acid sequences comprising Group 3, it is highly likely that these potential depolymerases are structurally similar.

We were unable to identify any coding sequences in the genomes of our isolated KLPN phages sharing high similarity to the four experimentally validated webervirus depolymerase sequences used to create our BLASTP database. Thus, it is likely that any depolymerase activity associated with the phages isolated in our study is due to enzyme(s) that remain to be characterized experimentally. As part of our previous analysis of the genome of phage KLPN1, we hypothesized that ORF34 and/or ORF35 may encode the depolymerase activity of phage KLPN1 as these sequences include a predicted endo-*N*-acetylneuraminidase/endosialidase domain (Hoyles et al., 2015). Further experimental work is required to determine whether these are functionally active depolymerases. As most of the plaques we observed had no discernible haloes, it may be that alternative mechanisms are used by weberviruses for penetrating the bacterial capsule. Depolymerase-independent penetration of the capsule by *Klebsiella* phages has been reported in the literature (Beamud et al., 2023).

### Genome-based analyses of publicly available phage data

A ViPTree proteome-based analysis of publicly available sequence data showed 330 genomes derived from isolated phages (*n*=265) and MAGs (*n*=65) belonged to the genus *Webervirus*, family *Drexlerviridae* (**Figure 1**). Our gene-sharing network analysis supported this finding (**Figure 2b**). Taxonomic assignment of phages using whole genome gene-sharing profiles has been shown to be highly accurate; a recent study showed that vConTACT2 produces near-identical replication of existing genus-level viral taxonomy assignments from the ICTV (Bin Jang et al., 2019). It has been suggested that genomes comprising a genus should be evaluated by phylogenetics with the use of ‘signature genes’ that are conserved throughout all members (Turner et al., 2021). Such analyses should always produce trees that are monophyletic. Using *terL* as a ‘signature gene’, we were able to show that the genus *Webervirus* is indeed monophyletic (**Figure 2a**). To assess the number of different species present within the genus, we used VIRIDIC to determine the intergenomic similarity between phage genomes (**Supplementary Figure D**; **Supplementary Table 4**). Guidelines suggest any two phages belong to the same species if they are more than 95 % identical across their entire genome (Turner et al., 2021). Genus-level separation occurs when phage genomes share <70 % nucleotide identity across their genome length (Turner et al., 2021). Based on these criteria, our results show that only weberviruses belonging to Cluster 1 represent species of *Webervirus sensu stricto*. Clusters 3–9, although identified as weberviruses using ViPTree and vConTACT2, do not represent species of *Webervirus*. The phage sequences associated with these clusters were derived from low-quality MAGs. As such, we recommend caution when using low-quality MAGs to determine taxonomic affiliations of *in silico*-generated phage sequences.

Cluster 2 phages were found to represent a novel genus (*Defiantjazzvirus*) of phage within the family *Drexlerviridae* (**Figure 1**, **Figure 2**, **Supplementary Tables 4 and 5**), with the genus *Defiantjazzvirus* most closely related to the genus *Webervirus*. All members of this novel genus reported to date infect a range of *Klebsiella* spp. (**Supplementary Table 2**).

As phages are among the most abundant biological entities on Earth, it is important to gain knowledge on their presence within different environments. We determined that weberviruses are distributed globally and predominated by phages associated with human faeces or water supplies contaminated with human faeces (**Figure 3**). Lack of detection in most of South America and Africa is likely due to absence of metagenomic datasets from these parts of the world rather than weberviruses not being represented in faecal samples from individuals living in countries within these regions. Compared with shotgun metagenomic datasets characterizing the total microbiota found in faeces, there are very few studies – worldwide – examining solely the intestinal virome, and PhageClouds is populated with phage genomes derived from virome datasets.

**Figure 3.**
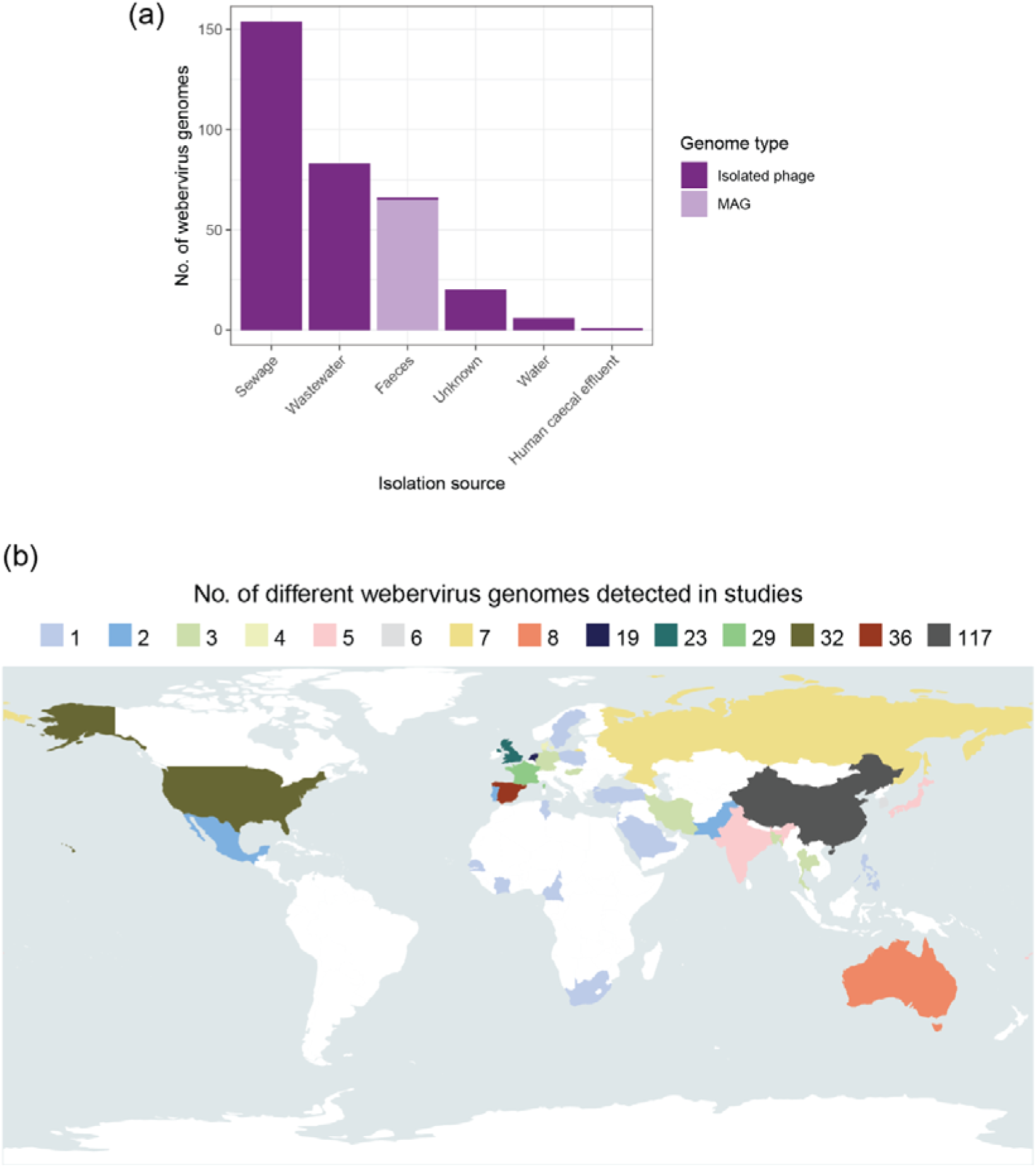
Distribution of weberviruses (a) Stacked bar graph showing the sources of the 330 webervirus genomes (*n*=265 isolated phages; *n*=65 MAGs). (b) Geographical distribution of 329 of the webervirus genomes included in this study (the location information was not available for one isolated phage, namely Klebsiella phage 5899STDY8049225).

**Figure 4.**
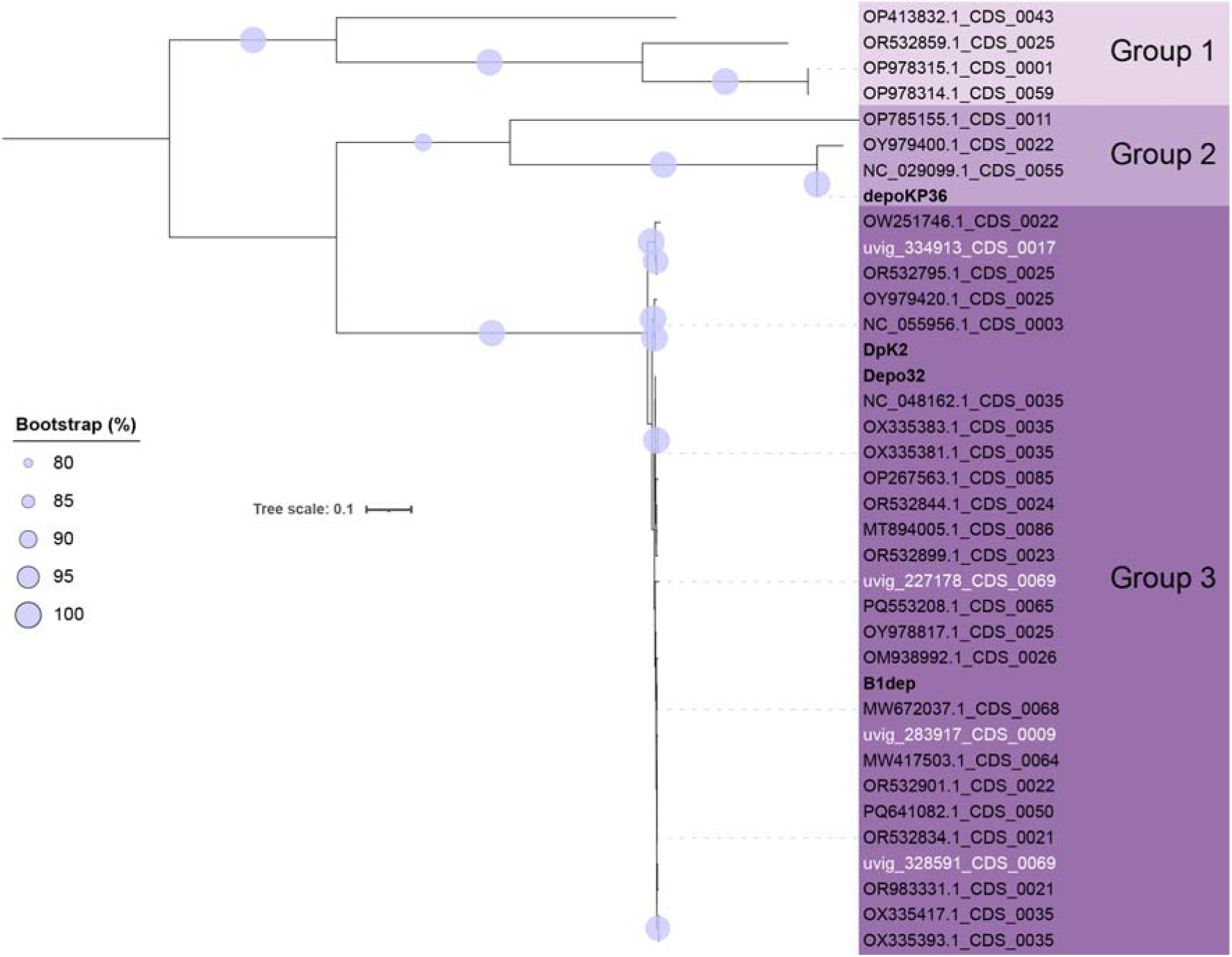
Phylogenetic analysis of depolymerases predicted to be encoded by weberviruses. The tree (maximum likelihood) is rooted at the midpoint. Bootstrap values are presented as a percentage of 100 replicates. Names of experimentally validated (i.e. functional) depolymerases are shown in bold black text; depolymerases predicted to be encoded by MAGs are shown in white text. Scale bar, mean number of amino acid substitutions per position

It was notable when curating our MAG dataset that none of the studies describing these data was able to predict hosts for the webervirus MAGs we have identified. Nor was the recently released tool iPHoP, specifically designed for use with MAGs (Roux et al., 2023). Our analysis using HostPhinder predicted webervirus MAGs infect *K. pneumoniae*. HostPhinder predicts the host species of a phage by searching for the most genetically similar phages in a database of reference phages with known hosts (Villarroel et al., 2016). Although the authors have shown that this whole-genome similarity-based approach is highly accurate, host range can be altered by a relatively low number of mutations, especially those localized to tail fibre proteins which are often determinants of host-cell specificity (Latka et al., 2021; Taslem Mourosi et al., 2022). In the present study, we used a strain of *K. variicola* to isolate three weberviruses and phages of this genus have also been isolated on *K. oxytoca* and *K. aerogenes*. Although it is highly likely that 329/330 weberviruses discussed herein are phages of *Klebsiella* spp., determination of host range via plaque assays is still informative, especially when determining therapeutic utility.

## SUMMARY

We successfully characterized seven novel weberviruses that infect clinically relevant MDR *Klebsiella* spp. We have trebled the number of authenticated webervirus genomes through combining genomic data from isolated phage and MAG datasets. In doing so, we have demonstrated the importance of interrogating MAG datasets to expand the availability of curated phage genome sequences for use in genomic and ecological studies, and highlighted the need to exercise caution when assigning low-quality MAGs to taxa.

## ACKNOWLEDGEMENTS

Consultant microbiologist Dr Frances Davies (Imperial College Healthcare NHS Trust) is thanked for providing access to clinical strains. We thank Emily Goren for providing mixed-liquor samples from Mogden Sewage Treatment Works (Thames Water). Dave Baker and the QIB core sequencing team are thanked for WGS library preparation and sequencing for the PS-prefixed bacterial genomes. Horst Neve is thanked for generating TEMs of phages vB_KpnS-KLPN2, vB_KpnS-KLPN3 and vB_KpnS-KLPN4.

LH, ALM and DN designed the study. PS, ALM, SG, TT and LH isolated and purified the phages. TCB and SJTD assembled and annotated the phage genomes. ALM and MK processed clinical isolates for whole-genome sequencing; PS and LH assembled and annotated the *Klebsiella* genomes. PS determined the antimicrobial profiles for the *Klebsiella* strains included in this study. FN determined the host ranges of the MAGs. SJTD did all bioinformatics work associated with the seven new phage genomes and the vConTACT2 analyses; LH did all other bioinformatics work associated with the MAGs. DN produced sequence data and TEM images. LH and ALM supervised PS. LH supervised TT, TCB and SJTD. LJH supervised MK. DN supervised SG. SJTD, DN and LH drafted the manuscript. All authors read and approved the final version of the manuscript.

## ETHICS

The study of anonymised clinical isolates beyond the diagnostic requirement was approved by an NHS research ethics committee (number 06/Q0406/20).

## Abbreviations

ICTV: International Committee on Taxonomy of Viruses
MAG: metagenome-assembled genome
MDR: multidrug-resistant
ST: sequence type
TEM: transmission electron micrograph
UHGG: Unified Human Gastrointestinal Genome.

## Data availability

The sequences for the seven new phage genomes described herein have been deposited in DDBJ/ENA/GenBank under accession numbers OM065837–OM065843. The genome sequences of bacteria described herein have been deposited under BioProject PRJNA917129. All supplementary material is available from https://figshare.com/projects/Weberviruses_are_gut-associated_phages_that_infect_Klebsiella_spp_/128516.

## Supplementary figures for

**Supplementary Figure A.**
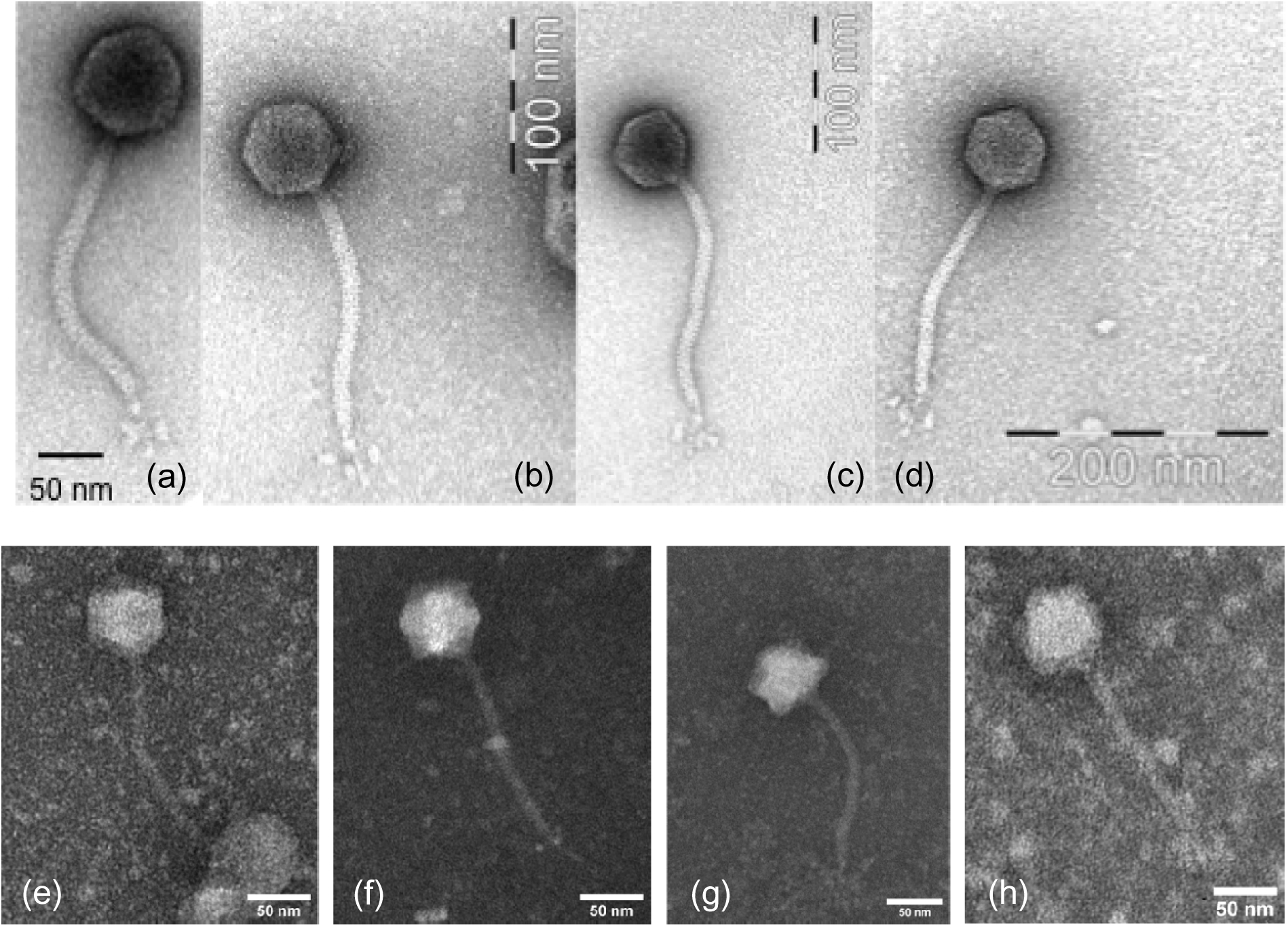
TEM images of (a) KLPN1, (b) vB_KpnS-KLPN2, (c) vB_KpnS-KLPN3, (d) vB_KpnS-KLPN4, (e) vB_KvaS-KLPN5, (f) vB_KvaS-KLPN6, (g) vB_KvaS-KLPN7 and (h) vB_KpnS-KLPN8.

**Supplementary Figure B.**
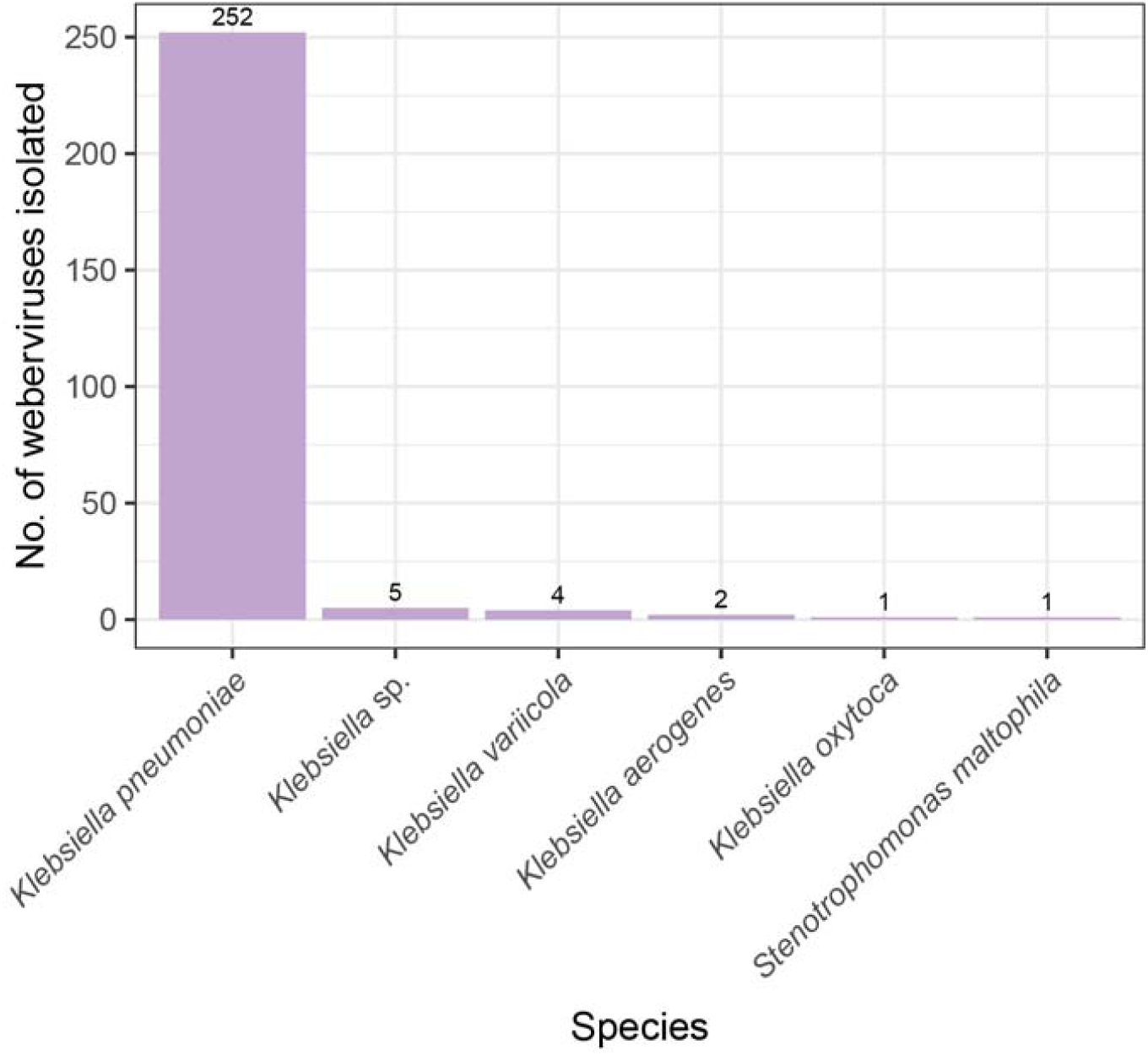
Weberviruses are most frequently isolated on *Klebsiella* spp., particularly *K. pneumoniae*. The isolation sources of the 265 isolate genomes (**Supplementary Table 2**) were collated and are summarized here.

**Supplementary Figure C.**
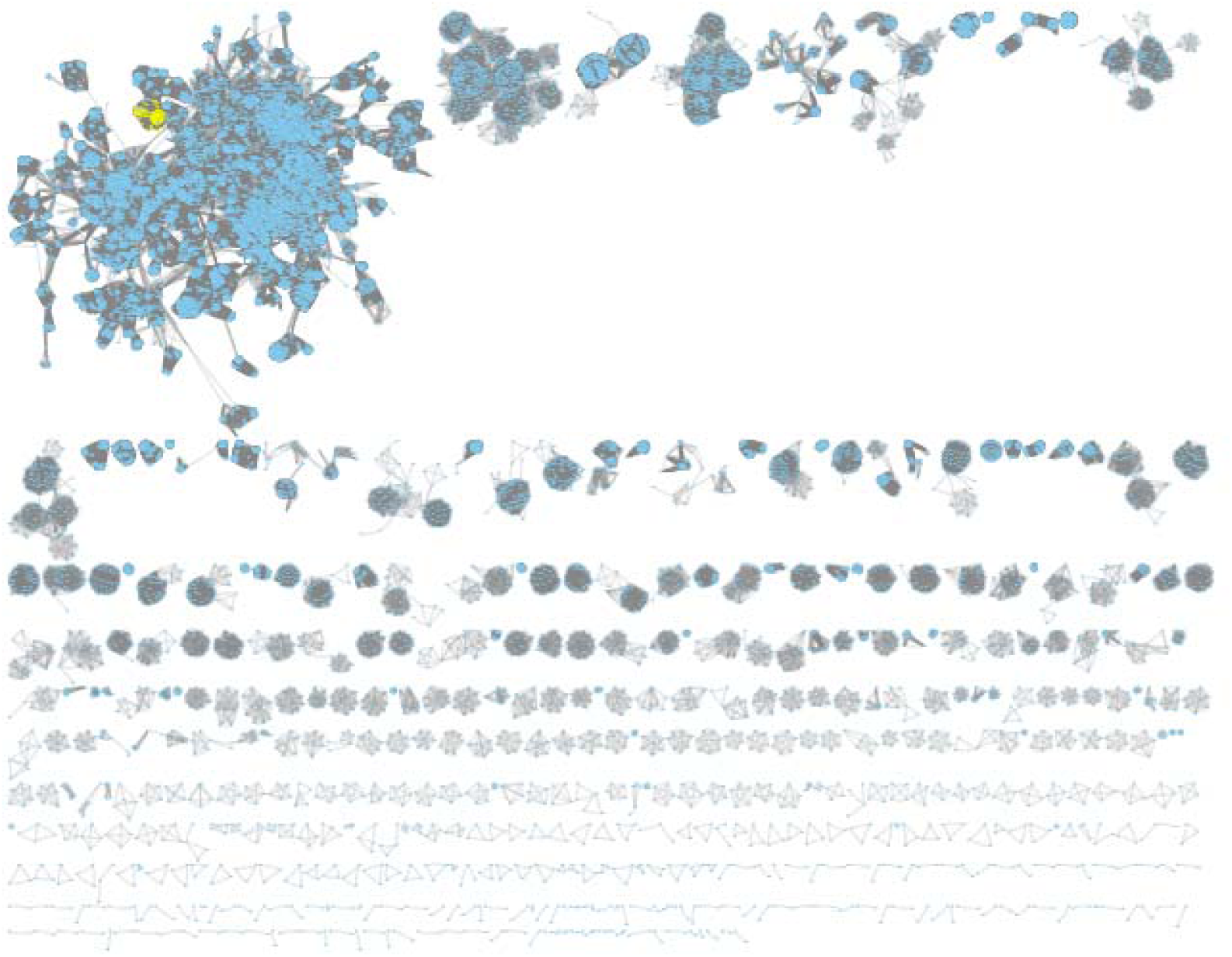
Full gene-based network generated using all genomes listed in **Supplementary Table 2** and **Supplementary Table 3**. Weberviruses are shown in yellow at the top left-hand side of the image. These and their first and second neighbours were identified and used to generate the subnetwork shown in Figure 2(b).

**Supplementary Figure D.**
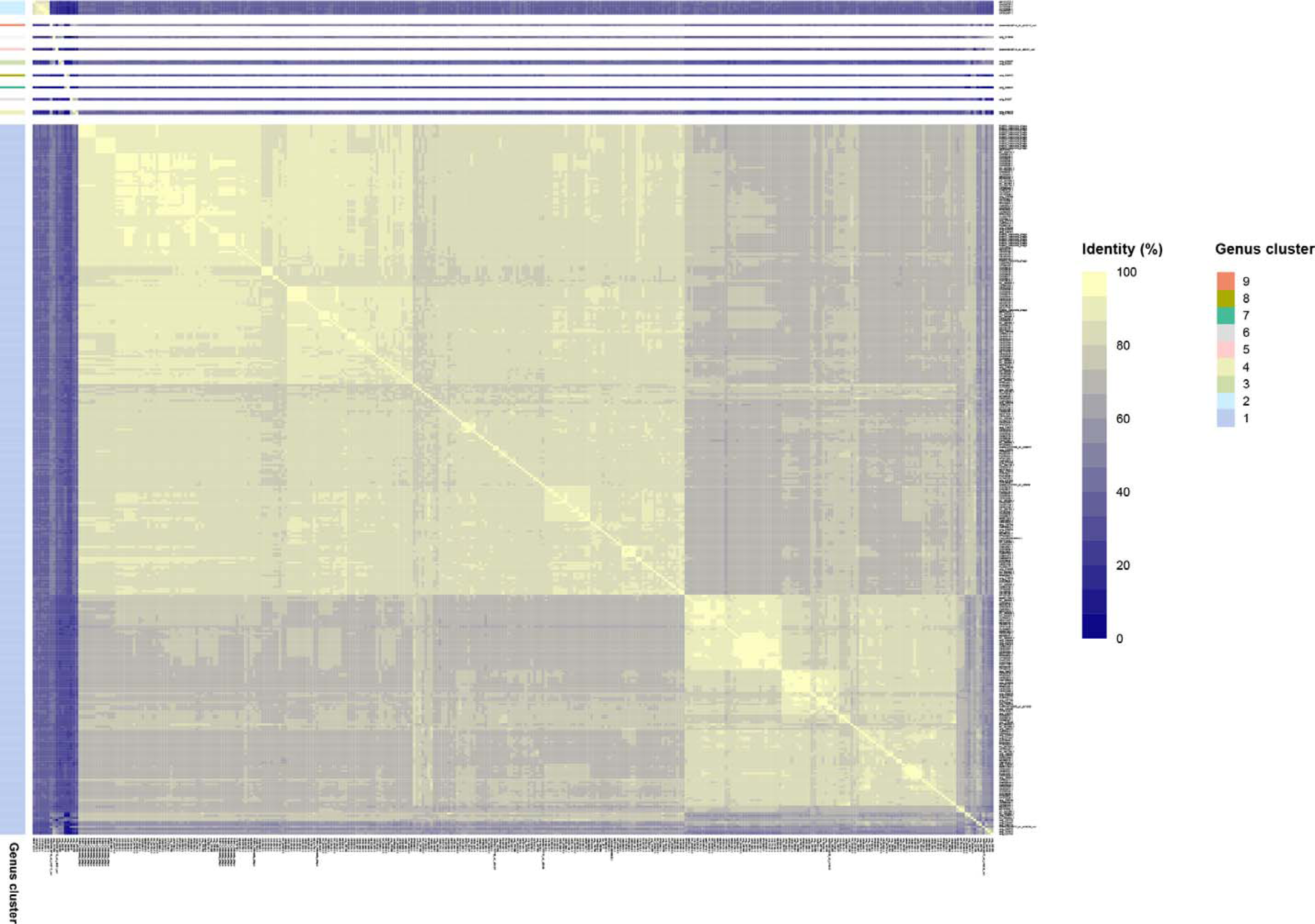
Heatmap showing bidirectional clustering of taxmyPHAGE similarity data for the 330 webervirus genomes and those of the novel genus *Defiantjazzvirus*. The numbered clusters correspond to the nine genus clusters detected by taxmyPHAGE with our curated set of sequence data.

**Supplementary Figure E.**
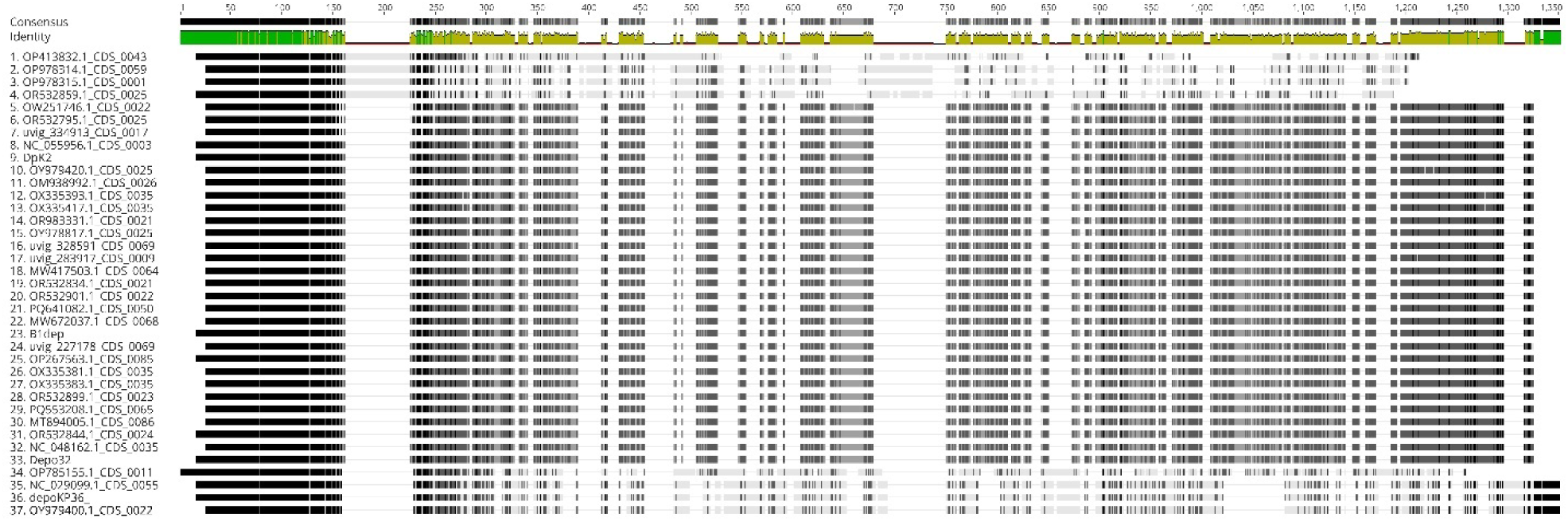
Amino acid alignment (Clustal Omega) of depolymerases identified in this study. A high level of sequence conservation (indicated in green) was identified at the N-terminal region of all the depolymerase sequences (first 100 amino acids).

